# Molecularly specific solubilization of therapeutic antibodies

**DOI:** 10.64898/2026.01.29.702671

**Authors:** Zexiang Han, Nadia A. Erkamp, Rob Scrutton, Giuseppe Licari, Olga Predeina, Andreas Evers, Pietro Sormanni, Tuomas P. J. Knowles

## Abstract

Understanding the effects of formulation excipients on protein solubility is a key part of physical chemistry and pharmaceutical sciences. While excipients are routinely employed to reduce the self-association of biologic drugs, their mechanisms of action remain poorly understood and are often assumed to be broadly nonspecific. Using a high-throughput combinatorial droplet microfluidic platform, we systematically survey and quantify how common pharmaceutical excipients affect the solubility of a diverse panel of therapeutic monoclonal antibodies (mAbs). We show that, while excipients are generally solubilizing, their effects vary substantially across different mAbs, with excipient-specific solubilization scores spanning dynamic ranges of approximately 7-fold to >200-fold across the antibody panel. Histidine, arginine and sodium chloride, in particular, engage in interactions characterized by unique molecular specificity, whereas sucrose effects are largely governed by nonspecific, solvent-mediated interactions. Correlating excipient performance with dynamical mAb molecular features from solvated full-length homology models allows us to dissect and quantify the relative contributions of molecular features governing excipient-mediated solubilization. We envision this new physicochemical understanding lays the groundwork for rational excipient selection and bespoke formulation design, with direct implications for accelerating protein therapeutic development for preclinical scenarios.

## Introduction

Antibodies have transformed the therapeutic landscape since the turn of the century, representing a potent class of biologic drugs capable of treating or mitigating a range of maladies from infections to cancers. Most antibody drugs approved by regulatory agencies like the U.S. Food and Drug Administration (FDA) are canonical forms of monoclonal antibodies (mAbs), praised for their high specificity, high potency and favorable safety profiles. Their commercial success is also evident from their substantial and still-growing market size, contributing to a global industry valued at over $200 billion.^1^

Two key and closely related developability traits of antibody-based therapeutics are colloidal stability and solubility.^2,3^ For antibody drugs intended for extensive patient populations or high-dosage regimens, high-yield production and stable formulation of antibodies are critical. Numerous commercial mAbs are formulated at ∼50–200 mg mL^−1^in liquid forms such as in vials or prefilled syringes to be amenable to subcutaneous injection, which enables self-administration and improves patient compliance. Proteins, however, become colloidally unstable at high concentration as short-range attractive interactions begin to dominate, resulting in opalescence and high solution viscosity.^4^These high-concentration formulations also experience various types of stresses including temperature changes or mechanical agitation, making proteins more susceptible to aggregation which inevitably leads to drug inactivation and/or aggregate-mediated immunogenicity. ^5^This constitutes a major challenge in biotechnology and the pharmaceutical industry. One strategy has been to engineer antibody mutational variants to reduce the intrinsic aggregation propensity of drug candidates.^3,6^Nonetheless, this strategy may impact functionality and can sometimes increase the polyreactivity of the drug.^7^

Excipients are inactive substances routinely added to biopharmaceutical formulations to fulfill functionally distinct roles. Essential components include buffers (e.g. histidine, acetate) for pH control and tonicity agents (e.g., NaCl, glycerol, sucrose) for osmolarity adjustment, while others, such as surfactants (e.g., polysorbates) or antimicrobial preservatives (e.g., metacresol), are added depending on the target product profile. Certain excipients, particularly amino acids, salts, and sugars, also function as colloidal stabilizers that improve antibody solubility and reduce self-association. Importantly, these stabilizers are not antibody-specific but modulate many proteins^8,9^and even non-proteinaceous colloids.^10^Many efforts, both experimental^11,12^and computational^13,14^, have been devoted to study antibody-excipient interactions in solution formats. However, these insights are often case- and context-specific^15–17^, limiting their generalizability across different mAb therapeutics.

Excipient formulations play a key role in biologic development.^18–22^The molecular basis of small molecule modulation of colloids such as therapeutic mAbs is largely unclarified, and machine learning approaches at the present stage are likely biased due to limited datasets. In particular, knowledge gaps persist regarding the efficacy of common additives in improving solubility and stability across clinically relevant antibodies. Conflicting evidence exists, such as regarding whether excipient binding is a good predictor of protein stabilization.^13,23^Therefore, a deeper understanding of antibody formulatability, beyond sequence-level variant optimization, is essential for biologics development.

In recent work, we introduced ultrahigh-throughput combinatorial droplet microfluidic platforms that map protein solubility and phase behavior across large chemical spaces.^24–26^Such methods dramatically reduce material consumption and measurement time compared to traditional assays, enabling high-resolution solubility phase diagrams over multiple dimensions. These methods have so far been used primarily to demonstrate general platform capabilities or to optimize solubility for individual proteins^25^, without systematically dissecting the molecularly specific effects of pharmaceutical excipients across a clinically relevant antibody landscape or quantitatively linking excipient responses to antibody-intrinsic physicochemical features.

Herein, we extend the platform to investigate whether common pharmaceutical excipients act the same or differentially on the solubility of a panel of therapeutic immunoglobulin G (IgG) antibodies, in the presence of histidine, sodium chloride, arginine, and sucrose. The high-resolution datasets enabled quantitative comparison of excipients in terms of their solubility-enhancing capabilities for a given antibody and across antibodies, providing a viable method to assess the formulatability of biologic drugs. Through *in silico* antibody structural modeling and physicochemical profiling, we establish physically interpretable correlations between excipient response and antibody molecular traits, thereby enabling bespoke formulation design for a given antibody based on sequence information alone. Our findings shed light on the intricate specificities of excipient-protein interactions and are expected to inform the rational design of biotherapeutic formulations as well as offer additional insights into the multifaceted roles small molecules play in protein-protein interactions.

## Results & discussion

### Measuring high-resolution solubility phase diagrams of therapeutic antibodies

PEG precipitation assay remains a common experimental technique to determine the relative solubility of proteins through molecular crowding and is routinely applied to examine the solution behavior of therapeutic antibodies.^27–29^Alternative assays, such as affinity-capture self-interaction nanoparticle spectroscopy (AC-SINS), albeit applicable for studying mAb self-association, are less suited for accurately quantifying excipient effects across excipient types.^30^We employed a combinatorial droplet microfluidics platform capable of generating thousands of compositionally distinct microdroplets within minutes, each encapsulating different concentrations of protein, PEG, and excipient(s) that serves as a unique chemical microenvironment for an aggregation reaction (**Figure 1**a).^24,25^This enables efficient and systematic screening of the chemical-concentration space with high throughput and reduced sample consumption. Using microfluidic PEG precipitation assay, we measured the relative solubilities of a panel of 14 IgG1 mAbs (**Supplementary File 1**), many of which are or based on marketed and/or clinical-stage drugs, in standard formulation buffer (10 mM histidine, pH 6.0). Notable features of this technique include an ultralow sample requirement, high-throughput data acquisition, and the ability to directly visualize mAb aggregates as subvisible particles.

**Figure 1.**
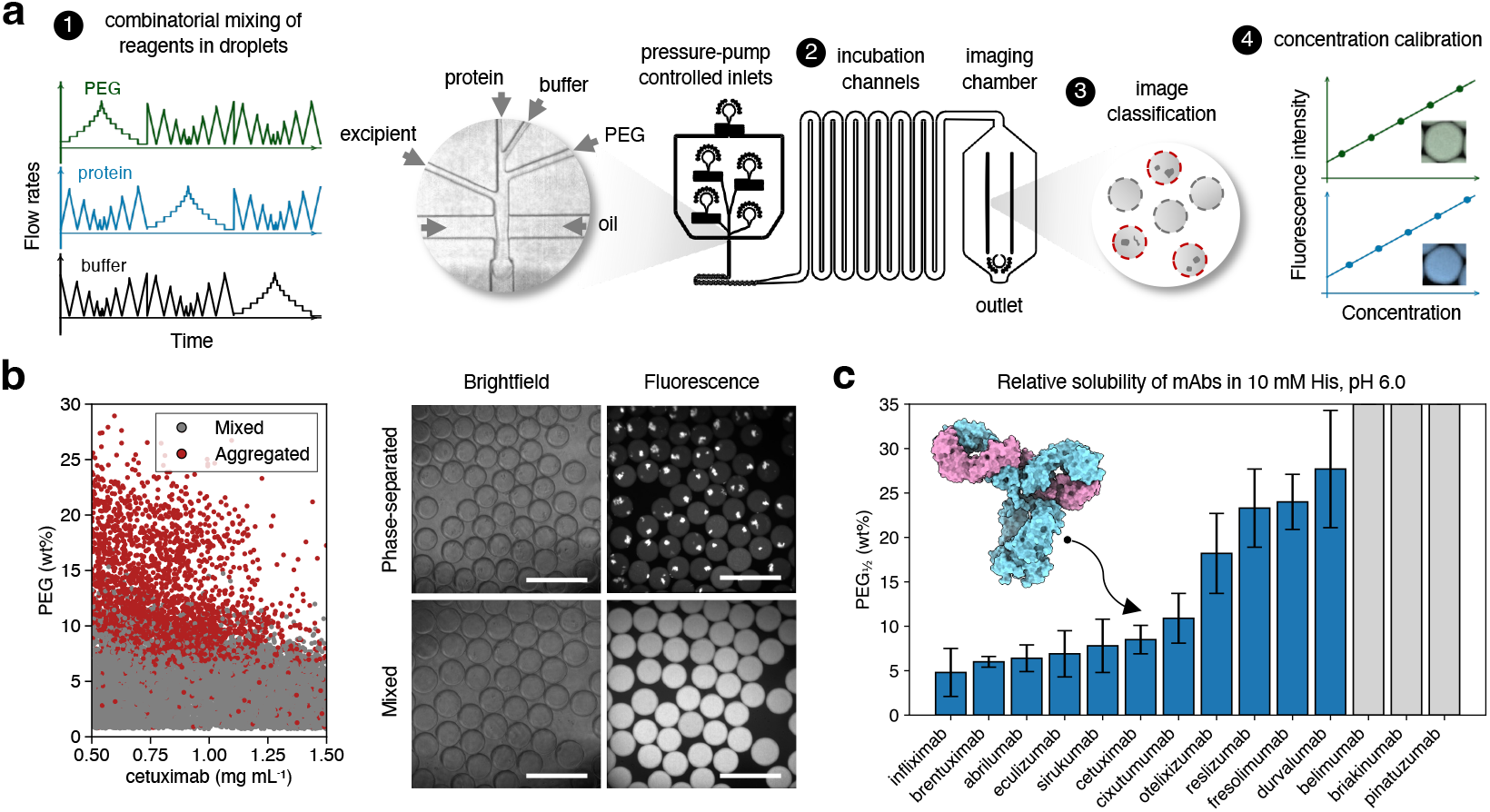
Combinatorial droplet microfluidics for evaluating the relative solubility of clinical antibodies. **a**, Overview of the microfluidic solubility screening assay workflow. (1) Combinatorial droplet generation using temporally defined flow profiles. (2) On-chip droplet incubation. (3) Droplet imaging and classification. (4) Concentration calibration for each ingredient. **b**, Solubility phase diagram of cetuximab around 1 mg mL^−1^using microfluidic PEG precipitation assay (number of data points *n* = 22,326). Shown on the right are representative images of microdroplets containing proteins (cetuximab) in the precipitated or fully mixed states. Scale bars = 200 *μ*m. **c**, Summary of the relative solubilities of therapeutic antibodies at 1 mg mL^−1^in standard formulation buffer. The error bars mark the 68% confidence interval as determined from the solubility profiles. Inset shows the full-length cetuximab structural model.

The droplet generation process is robust, and each experiment yields a uniform droplet population with a coefficient of variation below 5% (**Supplementary Figure S1**). Following microdroplet on-chip incubation, solubility classification was performed for identified droplets, totaling 22,326 data points for the experiment shown in **Figure 1**b whilst using only <50 *μ*g of the protein sample. Extrapolating the points of 50% aggregation probability that delineate the phase boundary, we determined the relative solubility of cetuximab at 1 mg mL^−1^to be PEG_1/2_ = 8.5±1.6 w/w% (**Supplementary Figure S2**). Owing to efficient mass transport within confined volumes of picoliter-droplets, increasing the incubation period beyond five minutes did not lead to significant variations in the measured relative protein solubility, indicating that near-equilibrium conditions were reached within our assay timeframe (**Supplementary Figure S3**).

**Figure 1**c summarizes the measured relative solubilities of a panel of clinically relevant mAbs, with their solubility profiles shown in **Supplementary Figure S4**. mAbs with a higher isoelectric point (pI) are generally more electrostatically stabilized in pH 6.0 histidine buffer (**Supplementary Figure S5**), consistent with previous reports.^31^Indeed, briakinumab, which because of its high pI has been associated with poor developability metrics such as poly-specificity^2^, did not precipitate out even for a PEG concentration as high as 30% under the same buffer condition. Given the typical trade-off between antibody binding specificity and self-association^7^, it could be more desirable to design suitable formulation recipes to prevent antibody aggregation at high concentrations without compromising their intended functional performance. 10 out of the initially considered 14 antibodies with a relative solubility below 25% PEG were selected for subsequent excipient formulation screening.

### Antibody solubilization by excipients

The selected panel of mAbs, all of which are post Phase-I in their development, represents a good set of candidates with confirmed developability for formulation excipient screening. Cetuximab, commercially known as Erbitux, is a marketed drug used to treat colorectal cancer and head and neck cancer.^32^We selected commonly used formulation additives in approved high-concentration antibody drug formulations, including histidine (His, as a modulator of buffer strength), sodium chloride (NaCl), arginine (Arg), and sucrose to study excipient effects on cetuximab. The concentration range of each excipient was chosen to closely align with those found in approved and marketed products.^9^By examining the phase behavior of fluorescently tagged antibody at a gradient of excipient concentrations, we obtained high-resolution datasets by which excipient performance can be quantitatively compared (**Figure 2**).

**Figure 2.**
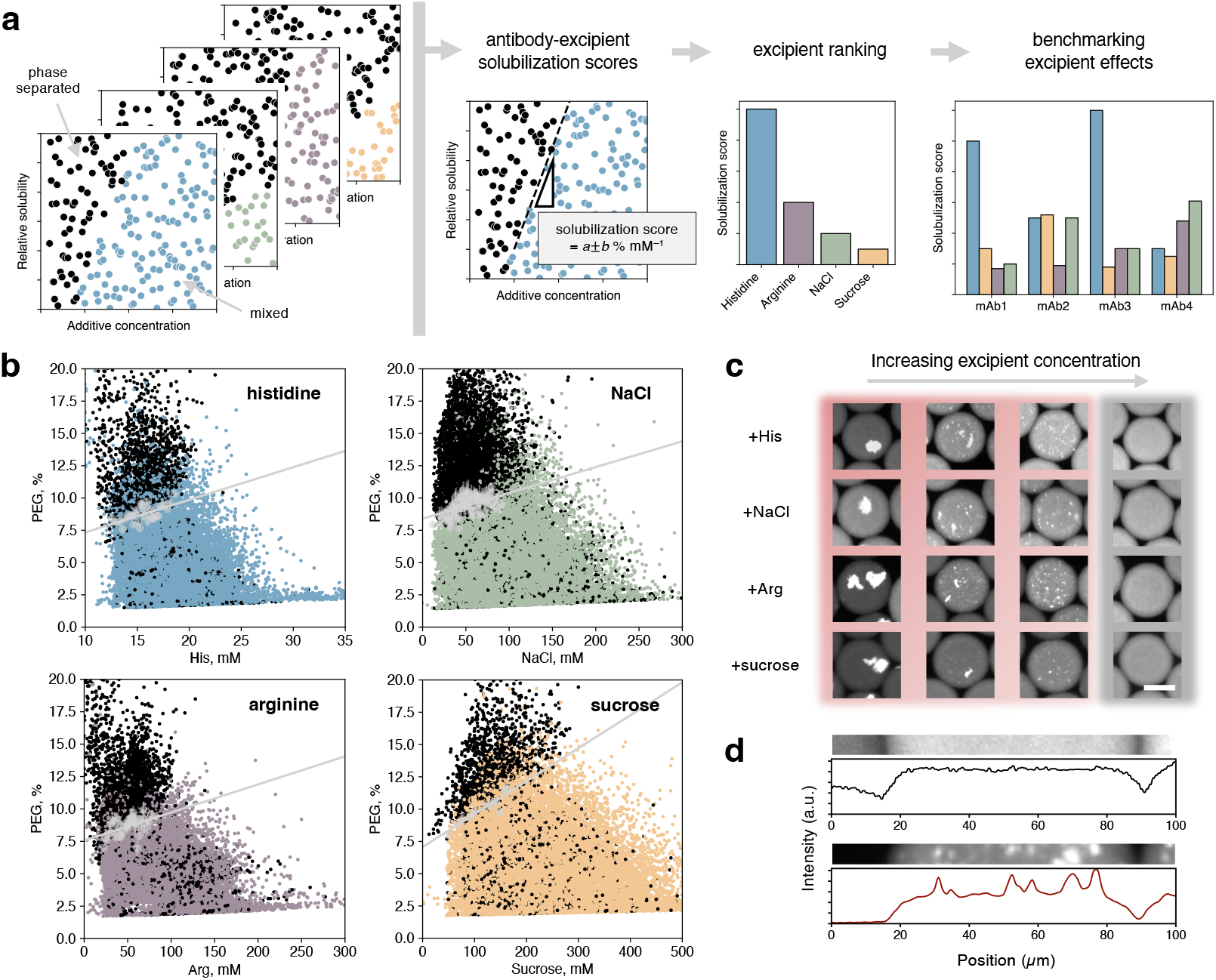
Formulation excipient analysis. **a**, Overview of excipient analysis using high-resolution phase diagrams. **b**, Solubility phase diagram of 1 mg mL^−1^cetuximab in the presence of common excipients: histidine (*n* = 16,291), sodium chloride (*n* = 21,424), arginine (*n* = 16,491), and sucrose (*n* = 21,702). **c**, Representative images of microdroplets showing antibody solubilization with increasing excipient input. Scale bar = 50 *μ*m. **d**, Fluorescent intensity line profiles for the mixed and phase-separated states.

**Figure 2**b illustrates how the relative solubility of 1 mg mL^−1^cetuximab varies with the addition of four common excipients. The % PEG crowder needed to induce mAb precipitation shifted to higher values with increasing excipient concentration, *i.e*., excipient addition led to an increase in the relative solubility of the antibody for all four compounds. Our microfluidic screening assay takes advantage of image analysis, allowing for the direct visualization of mAb aggregates, specifically subvisible particles, a feature that is lacking in ensemble measurements. No excipient-specific effects on the mAb aggregate morphology were seen at the micrometer length-scale. Upon PEG precipitation, the antibody exhibited two common precipitated states (**Figure 2**c). In conditions with no excipients (**Figure 1**b) or at low concentrations, large, amorphous aggregates were present, featuring irregular and relatively compact structures. In contrast, at higher excipient concentrations, the antibody either formed smaller speckles with diameters of 1–5 *μ*m or remained in fully mixed state. This range of morphologies reflects the varying phase behavior of the protein across the different conditions surveyed. The precipitated states are protein-rich in nature as they deplete fluorescently labeled mAb molecules from the neighboring environment within microdroplet reaction chambers (**Figure 2**d).

We applied linear regression analysis to the solubility phase boundary, and the obtained slope was used as a normalized metric for evaluating the excipient’s solubilizing efficiency, generating a *solubilization score* for each antibody-excipient pair. A steeper phase boundary slope corresponds to a higher solubilization score, meaning that the same solubility enhancement can be achieved with a lower molar concentration of excipient addition. To ensure that the solubilization score was not biased by fluctuations in individual fits, we inferred the score using a binning strategy, and slope estimates were obtained across a wide range of minimum-cluster thresholds. The resulting distribution showed a consistent central value, indicating that the solubilization score can be robustly determined (**Supplementary Figure S6**). For cetuximab, the excipients in order of decreasing solubilizing efficiencies are His (0.266 ± 0.041 % mM^−1^), Arg (0.034 ± 0.003 % mM^−1^), followed by sucrose (0.024 ± 0.003 % mM^−1^) and NaCl (0.023 ± 0.003 % mM^−1^) performing similarly well.

### Excipient responses can show high variability and molecular specificity

The mechanisms underlying the influence of excipients on protein colloidal stability and solubility remain poorly understood, owing to both the envelope of molecular interactions at play and their dynamic nature. Indeed, biologic formulation design frequently necessitates extensive trial-and-error, or rely heavily on previously approved formulation recipes. To understand excipient-mediated solubilization of clinically relevant antibodies, we established a panel-wide reference by comparing experimentally determined solubilization scores across therapeutic antibodies (**Figure 3**a).

**Figure 3.**
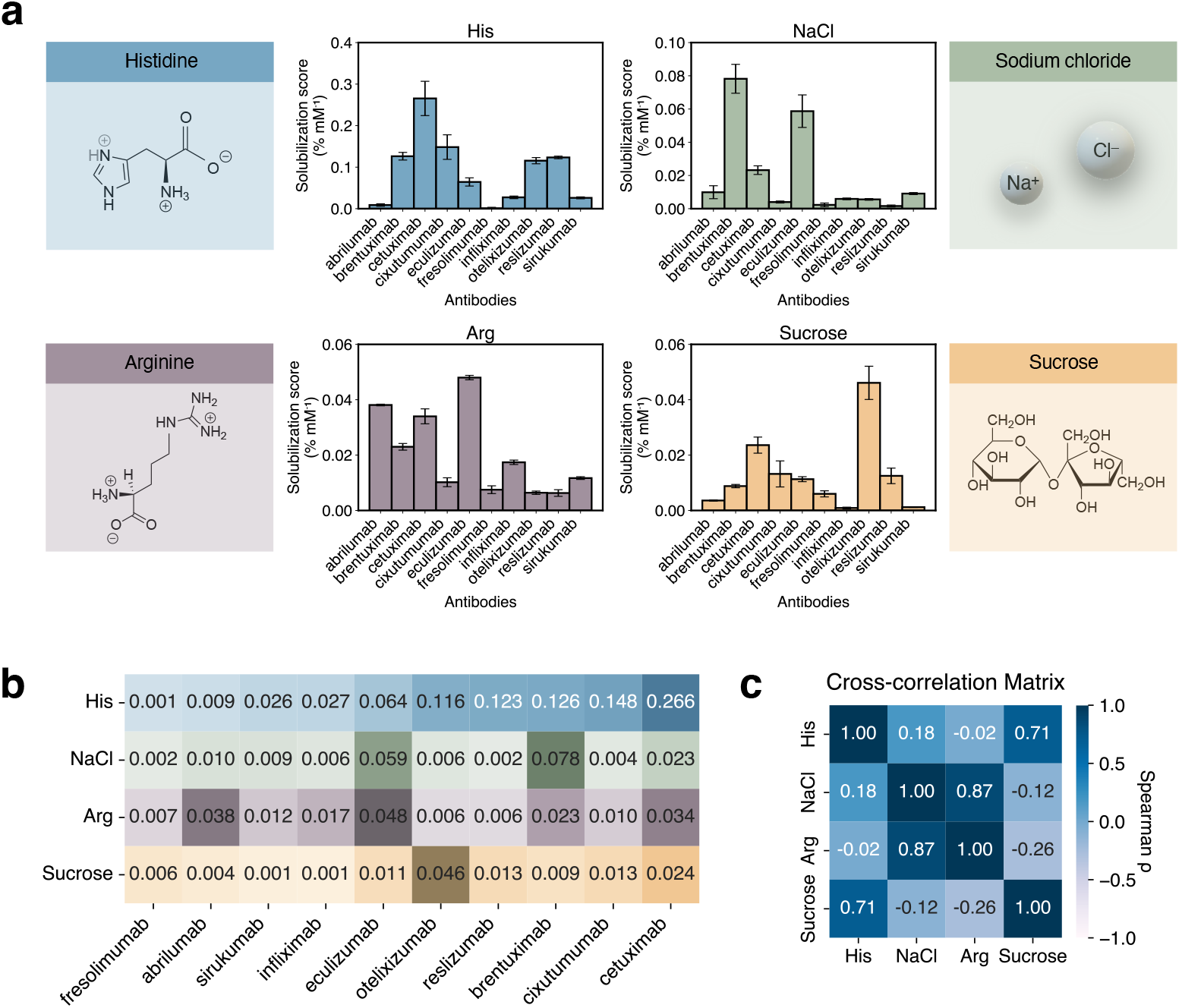
Evaluating solubilization efficiencies of common formulation excipients. **a**, Global comparison of solubilization scores for all protein-excipient pairs studied. **b**, Heatmap summarizing the data in (a). **c**, Cross-correlation matrix of the excipient solubilization scores based on Spearman’s ρ.

With a normalized heatmap shown in **Figure 3**b, our analysis reveals several key findings regarding the role of excipient additives on protein solubility. Firstly, the solubility-enhancing effects of excipients are often highly antibody-specific. A particularly striking disparity was observed for the histidine buffer: while histidine addition enhanced the solubility of cetuximab significantly, the same conferred minimal benefit to fresolimumab, with a greater than 200-fold difference in solubilization scores. This substantial disparity underscores that the effects of small molecule additives, including those used as the buffer, are not universally applicable across antibody drugs. Secondly, for any given protein, the efficacy of different excipients can vary substantially, as exemplified by almost all mAbs studied here. Finally, no single excipient universally maximized solubility across the tested antibody panel, indicating that a one-size-fits-all formulation condition for antibody drugs is unlikely. This highlights the need for targeted, rational excipient selection to optimize solubility for the therapeutic protein in question. To contextualize these findings, we examined the solubilization efficiency of the excipients on globular protein bovine serum albumin (BSA), which is smaller and has a simpler surface structure compared to mAbs. We hypothesize that fewer excipient molecules are needed to interact with and stabilize a single BSA molecule compared to an IgG, resulting in higher solubilization scores for BSA. This is indeed observed for all four excipients, albeit to varying extents (**Supplementary Figure S7**).

Because NaCl, Arg, and sucrose were titrated in a constant 10 mM histidine buffer, we assessed whether excipient responses merely track the histidine baseline across antibodies by examining the cross-correlations of solubilization scores (**Figure 3**d and **Supplementary Figure S8**). The scores of NaCl and Arg showed negligible correlation with histidine, indicating their mAb-specific effects are primary and not artifacts of the histidine background. On the contrary, sucrose solubilization score exhibited a strong correlation with those from histidine and a relatively small dynamic range across the mAb panel. First, sucrose’s effect is overwhelmingly nonspecific and uniform across mAbs, as shown by its small dynamic range compared to histidine. Second, and more critically, sucrose is a well-established excipient that operates *via* preferential exclusion and the strengthening of the hydrophobic effect.^33–35^We propose that sucrose does not mimic histidine’s specific interactions but rather globally reinforces the protein conformation stabilized by them. Therefore, the strong correlation could reflect that the magnitude of sucrose’s nonspecific stabilization is dependent on, and amplifies, the degree of specific stabilization first provided by histidine. Notably, specific binding interaction of sucrose with antibodies, such as BMSmAb^12^, has been documented in case reports; nevertheless, these findings appear rare and antibody-specific. Our data, along with the observation that sucrose increased thermal unfolding transition temperatures (**Supplementary Figure S9**), supports and reaffirms a broader, nonspecific mechanism of preferential exclusion by which sugar molecules stabilize therapeutic antibodies.

We additionally note a strong correlation between the solubilization scores of Arg and NaCl (**Supplementary Figure S8**). However, this co-variation reflects similarities in their macroscopic solubility profiles under our assay conditions and does not imply identical underlying mechanisms, as the two excipients differentially modulate antibody thermostability (**Supplementary Figure S9**) and are known to engage distinct physicochemical interactions (see discussion later).

### Unraveling physicochemical rules dictating excipient responses

Our analysis thus far highlights that excipients do not exhibit a uniform effect on all therapeutic mAbs. Additionally, the comparison between solubilizations of BSA and IgG suggests molecular determinants of excipient responses. We then set out to understand the observed heterogeneity in the solubilization scores between antibody-excipient pairs by examining the physicochemical characteristics of mAbs. Full-length amino acid sequences were used to compute sequence-level properties, and structural properties were computed based on atomistic molecular dynamics (MD) simulations of homology-based IgG molecular models to approximate their conformational dynamics in solution (**Figure 4**a).^36^Specifically, some size-related and patchiness metrics were excluded as they were largely invariant across the antibody panel, reflecting the highly conserved IgG1 architecture and the limited simulation time employed (**Supplementary Figure S10**). We retained only 19 features for which typical within-antibody fluctuations were small relative to the panel-wide spread, producing a set of sufficiently differentiating features for analysis (**Figure 4**a).

**Figure 4.**
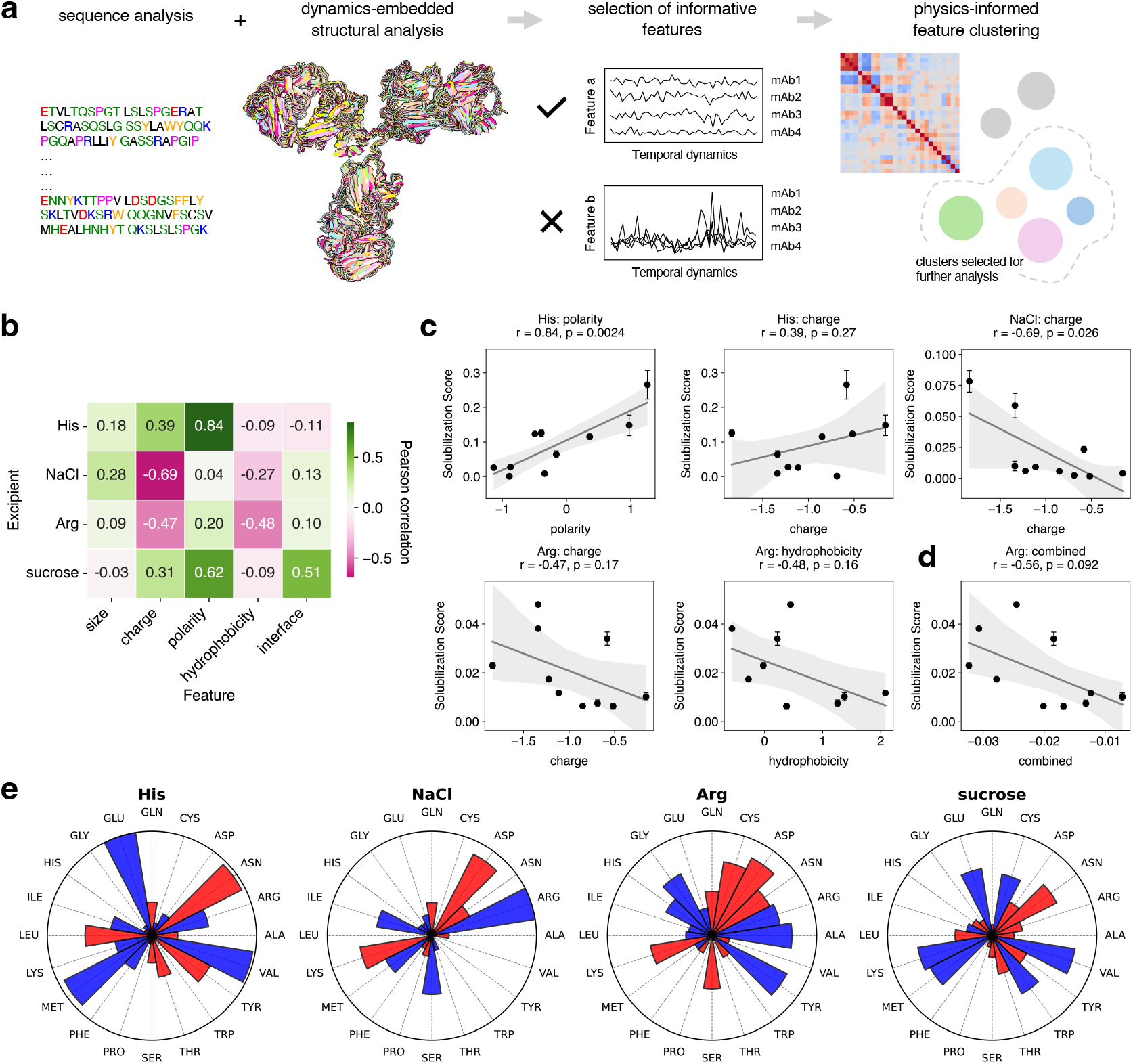
Understanding specific excipient effects on antibody solubility through intrinsic physicochemical profiles. **a**, Physics-based clustering of mAb physicochemical properties using full-length IgG sequence and structural models. Antibody visualization was performed using ChimeraX.^43^**b**, Heatmap of Pearson correlation coefficients on excipient-specific solubilization scores against feature clusters. **c**, Scatterplots showing the prominent correlation(s) for excipients His, NaCl and Arg, with linear regression fits and 95% confidence intervals as error bands. Pearson correlation coefficients (r) and associated p-values are indicated for each feature. **d**, Scatterplot showing Arg solubilization score plotted against a combined score derived from multiple linear regression of both charge (relative weighting = 0.497) and hydrophobicity (relative weighting = 0.503). **e**, Residue-level analysis. Here, the relative contributions of individual amino acid residues to the solubilization behavior of mAbs are shown for each excipient. Each polar sector represents one amino acid, and the radial length of the sector corresponds to the normalized magnitude of correlation between residue-specific solvent-accessible surface area (SASA) and the solubilization score for ease of visualization (red = positive, blue = negative). The plots reveal which residue types most strongly modulate antibody solubility in the presence of different excipients, highlighting distinct physicochemical signatures across His, NaCl, and Arg modulations.

These features were clustered into five interpretable property categories: size and hydrodynamics, charge, polarity, hydrophobicity, and light-chain/heavy-chain (LC/HC) interfacial properties. Composite z-scores derived from these clusters were subsequently correlated with experimentally measured solubilization scores, revealing molecularly specific stabilization effects of different excipients (**Figure 4**b).

Histidine is a commonly used amino acid buffer for IgG formulations showing a dual mechanism of action at pH 6, where the zwitterionic His^0^and protonated His^+^species co-exist.^37^This could in principle enable the simultaneous contribution of electrostatic screening and imidazole-mediated interactions. While our results were consistent with this interpretation, we found a dominant role of polarity in dictating mAbs’ response to histidine, as measured by dipole moment and polar solvent-accessible surface area (SASA) (**Figure 4**c and **Supplementary Figure S11**). Supporting this, residue-level analysis pinpoints tyrosine and asparagine residues of mAbs as potential interaction sites, where mAbs with more exposed polar Asn residues benefit the use of histidine for stabilization and solubilization. The added histidine molecules likely interact via π-π, hydrogen-bonding, and hydrogen-π interactions through the polar imidazole heterocycle to mediate its effects (**Figure 4**e).^38,39^Critically, the absence of any significant correlation with hydrophobicity-related metrics powerfully argues against a role for hydrophobic interactions *via* the imidazole ring. This indicates that His-mediated stabilization of therapeutic mAbs is fundamentally polar and, to a lesser degree, electrostatic in nature (**Supplementary Figure S11**), rather than driven by the hydrophobic effect.

Sodium chloride, in stark contrast to the specific interaction of His, operates through nonspecific electrostatic screening. Its efficacy to solubilize mAbs is uniquely associated with the suppression of electrostatic interactions, as evidenced by strong positive correlations with charge metrics. We further note that the investigated antibody panel is biased toward lower-pI species, and the composite charge z-scores are predominantly negative. The observed correlations reflect relative differences in charge within this restricted range rather than the presence of net positively charged antibodies. Therefore, NaCl excels at dampening the long-range electrostatics driving the self-association of problematic antibodies and provides diminishing returns for highly positively charged antibodies that are already electrostatically stabilized.^18,40^

Arginine presents another distinct and specialized mechanism of mAb solubilization. Arg molecules possess a guanidinium group that is positively charged in solution, and in line of this, our analyses identified the nearly-equal contribution of electrostatics and protein hydrophobicity in Arg-mediated stabilization (**Figure 4**c and d). The guanidinium group of Arg is uniquely suited to form hydrogen bonds with charged regions in antibodies, thereby neutralizing the long-range electrostatic interactions that can drive mAb self-association. On the other hand, the inverse correlation between Arg’s solubilization score and hydrophobic surface area reflects a trade-off in its mechanism. While Arg’s guanidinium group provides potent electrostatic stabilization, its chaotropic character concurrently disrupts hydrophobic effect and weakens cation-π interactions.^41,42^In mAbs with larger hydrophobic surfaces, this destabilizing effect offsets a greater portion of the electrostatic benefit, reducing the net solubilization efficiency per Arg added. Specifically, we found Arg efficacy to depend on hydrophobic SASA, with both alanine and tyrosine being likely interaction sites (**Figure 4**e and **Supplementary Figure S11**). Since the clinical-stage mAbs investigated have good developability properties, their intrinsic stability could buffer the magnitude of this mild destabilization, as separately seen in the reduced thermostability (**Supplementary Figure S9**).

Finally, sucrose solubilization score, as measured in histidine buffer, shows a strong correlation with the score for histidine and their broader correlation profiles share multiple similarities, consistent with its nonspecific stabilization behavior. In fact, the sucrose shows a moderately positive dependency on interfacial stability between the light and heavy chains (**Figure 4**b and **Supplementary Figure S11**). This is in agreement with the view that preferential exclusion scales with native state stability, making it easier for sucrose to stabilize mAbs with higher intrinsic interfacial stability.^33^

Overall, we have demonstrated the effects of common excipients on protein solubility are not promiscuous, even for a system as conserved in molecular attributes as mAbs. Despite the inherent structural heterogeneity and conformational plasticity of mAbs in solution, we have uncovered physically interpretable correlations between mAb molecular properties and their excipient response profiles. These correlations provide mechanistic insight into the dominant interaction modes engaged by different excipients and quantify how competing physicochemical interactions contribute to solubilization. Because the excipients are used at much higher concentrations compared to the antibodies, we hypothesize that the excipients form a protein-feature-dependent solvation shell around the proteins, mediating colloidal stabilization.^10^

Our work thus complements standard developability studies by shifting the focus from intrinsic aggregation tendencies to rational, mechanism-informed formulation design. Because the studied mAbs are representative of clinically relevant IgG1 antibodies^22,36^, these findings are likely generalizable across many therapeutic antibodies, advancing formulation from a largely empirical process toward a more predictive framework.

## Conclusions

In this work, we employed a microfluidic PEG precipitation assay for solubility measurements of clinically relevant antibodies and systematic formulation excipient profiling. A key practical advantage of our approach is the collection of high-resolution phase diagrams using minimal amounts of proteins, overcoming the material and time constraints of conventional solubility assays. Leveraging the high-resolution datasets, we introduced the solubilization score as a transferable metric for quantifying and systematically comparing excipient efficacy across proteins. Our data enabled excipient ranking for one IgG and demonstrated that, for a single excipient, its performance is highly variable between IgGs, indicating that formulation additives do not exert universal actions. By integrating phase stability profiles with *in silico* dynamics-embedded modeling, we revealed the molecular grammar underlying excipient-mediated solubilization and antibody-excipient interactions, providing mechanistic interpretations. We established that histidine, arginine, and sodium chloride, although all charged in solution, are not interchangeable but constitute a toolkit of distinct mechanisms, with excipients exerting differential efficacies depending on specific molecular features of the antibody. Together, these results move beyond a purely empirical paradigm of formulation by providing a mechanistic, physics-based framework that can support the rational design and optimization of antibody formulations. More broadly, we present a generalizable strategy for quantitatively dissecting small molecule-protein interactions, with implications for advancing the physical chemistry of protein and colloidal solutions across biotechnology and medicine.

## Methods

### Materials and antibodies

Polyethylene glycol (PEG, average molecular weight Mw 6000 g mol^−1^), sodium chloride (NaCl), L-histidine, L-arginine, and sucrose were purchased from Sigma Aldrich.

IgG1 antibodies were sourced from different pharmaceutical companies or expressed and purified by GenScript. Belimumab, briakinumab, cetuximab, cixutumumab, durvalumab, otelixizumab, and pinatuzumab were provided by the Merck Group. Brentuximab and sirukumab were provided by Novo Nordisk. Infliximab was purchased from Cambridge Bioscience Ltd. Abrilumab, eculizumab, fresolimumab, and reslizumab, for which known fragment-variable Fv regions were grafted onto a common IgG1 Fc domain, were acquired from GenScript. Mouse IgG1 was purchased from Thermo Fisher Scientific. The as-received antibodies were purified by size exclusion chromatography (Superdex 200 Increase 10/300 GL column) in 10 mM histidine, pH 6.0, with 150 mM NaCl and buffer-exchanged into 10 mM histidine prior to use.

### Preparation of microfluidic devices

Polydimethylsiloxane (PDMS)-based microfluidic devices were prepared following the standard soft lithographic protocol. PDMS prepolymer (Sylgard 184 Silicone Elastomer Kit) was mixed with the curing agent in a 10:1 w/w ratio and poured onto the developed silicon master. Degassed PDMS was then cured at 65 °C for 2 h. Upon demolding the PDMS slab, inlets and outlets were punched in using 0.75-mm biopsy punch. Cleaned PDMS was bonded to a glass slide *via* oxygen plasma treatment. The devices were subsequently annealed on a hot plate at 95 °C for 5 min, and the microchannel surface was hydrophobically treated with 2% trichloro(1H,1H,2H,2H-perfluorooctyl)silane (Sigma Aldrich) in HFE-7500 engineering fluid (Novec^™^, 3M). Excess oil was removed with the help of nitrogen stream before further annealing the device at 95 °C for 10 min.

### Combinatorial droplet microfluidics

A custom-built microfluidic platform was used to combinatorially prepare, incubate and image the microdroplets for solubility screening.^24^The microfluidic device consists of five inlets, a flow-focusing junction, a series of microchannels, and an imaging chamber. Antibodies were fluorescently labeled with Alexa Fluor 488 succinimidyl ester (Thermo Fisher Scientific). PEG solution and excipient solutions were freshly prepared in histidine buffer with appropriate pH adjustment and barcoded with 5 *μ*M Alexa Fluor 647 carboxylic acid and Alexa Fluor 546 carboxylic acid, respectively. Pressure-controlled pumps (OxyGEN, Fluigent) were used to temporally define the flow rates of HFE-7500 oil containing 2% fluorosurfactant (RAN Biotechnologies), antibodies, PEG, buffer, and excipient solutions into the device. Designing suitable flow profiles for each aqueous component enables the combinatorial mixing of PEG, protein, and additives within microdroplets. The oil phase flow rate was kept around 50 μL h^−1^and the sum of all aqueous flow rates at 105 μL h^−1^. Water-in-oil droplets were generated and confined between two parallel planes, resulting in a discoid shape due to physical compression, with a typical apparent diameter of 80–100 *μ*m. Droplets were incubated for approximately 5 min by flowing through serpentine microchannels to facilitate reagent mixing and continuously imaged using an epifluorescence microscope upon excitations at 488 nm, 561 nm, and 630 nm, respectively.

### Image analysis

Acquired images were analyzed similar to previously reported.^24^Briefly, droplets were identified with a circularity filter and subsequently classified as soluble or aggregate-containing using a pretrained convolutional neural network. Images on phase-separated proteins, including condensates, small speckle assemblies, and large, irregular aggregates, were used for data training. For every experiment, a calibration curve of fluorescent intensities *versus* known concentrations for each barcoded component was constructed, and the concentrations of antibodies, PEG and additives were subsequently determined. This enabled the construction of a solubility phase diagram.

### Calculation of the solubility profile

For each phase diagram, a probability map was generated using a support-vector machine algorithm. The solubility profile for each antibody plots the mAb aggregation probability as a function of PEG concentration at a fixed protein concentration (*i.e*., 1.0 mg/mL^−1^). A sigmoidal function of the form *P*(*x*) = (1 + exp(−*k*(*x*−*x*_0_)))^−1^ was fitted using nonlinear least squares regression to the aggregation probability data, where *x* represents the PEG concentration, *x*_*0*_ is the midpoint, and *k* describes the steepness of the curve. Relative solubility was reported as the PEG concentration at 50% aggregation probability, with uncertainty defined as the interval between 16% and 84% probabilities.

### Calculation of the solubilization score

The solubilization score for each antibody-excipient pair was obtained from high-resolution single-droplet phase diagrams as the slope of the inferred phase boundary using a binning-based classification procedure. Adaptive bins were constructed such that each bin contained at least a specified minimum number (minN) of both phase-separated and mixed droplets. This aggregation stabilized the local classification problem by ensuring that each bin contained sufficient information from both classes using balanced accuracy. The PEG value yielding the highest balanced accuracy was assigned as the local phase boundary, and bins with accuracy < 0.75 were excluded. Average phase-boundary slope and corresponding standard deviation were computed over minN = 30–100.

### Thermostability measurements

The thermal stability of antibodies was evaluated via intrinsic fluorescence measurements using the NanoTemper Tycho NT.6 system. Proteins were formulated at 1 mg mL^−1^in 10 mM His buffer at pH 6.0, and separately, with the addition of excipients. Samples were loaded into Tycho glass capillaries and subjected to a heating protocol from 35°C to 95°C at a rate of 0.5°C s^−1^. The first derivative of the resulting fluorescence ratio curve was calculated. The onset temperature of the thermal denaturation T_onset_ was calculated as the temperature at which the first derivative showed detectable deviation from the baseline with threshold set to 10^−3^. Measurements were performed in duplicates.

### Modeling full-length antibodies

A total of 65 clinical-stage antibodies were considered, including those investigated experimentally in this work. The full-length antibody sequences are provided in **Supplementary File 1**. We constructed full-length antibody homology models using the Antibody Modeler module of Molecular Operating Environment (MOE) 2022.02.^44^The Amber10:EHT force field was employed in combination with the Generalized Born implicit solvent model^45^, with internal and external dielectric constants set to 4 and 80, respectively. Nonbonded interactions were smoothly switched off between 10 and 12 Å. All N- and C-termini were retained in their standard, charged forms. Following model generation, structures were prepared in MOE to relieve steric clashes and to assign protonation states at pH 6.0 using the Protonate3D protocol.^46^Final models were energy minimized until the energy gradient was below 10^−2^kcal/mol·Å^2^.

### Molecular dynamics simulations

For classical atomistic molecular dynamics (MD) simulations, the homology models were embedded in a rectangular water box with at least 20 Å between any protein atom and the box boundary, and neutralized with NaCl using Visual Molecular Dynamics (VMD).^47^Equilibration runs were performed with Nanoscale Molecular Dynamics (NAMD) 2.14, whereas production simulations were carried out with NAMD 3.0b6.^48^The CHARMM36m force field^49^was used to describe the protein, and TIP3P water^50^was used for solvation. Nonbonded interactions were computed with a 12 Å cutoff and a switching function starting at 10 Å. Long-range electrostatics were treated using the Particle Mesh Ewald (PME) method,^51^with a grid spacing of 1 Å^−3^. All bonds involving hydrogen atoms were constrained using the SHAKE algorithm.^52^Pressure was maintained at 1 atm via a modified Nosé–Hoover Langevin piston,^53^and the temperature was controlled at 300 K with a Langevin thermostat. A timestep of 2 fs was used, evaluating Lennard-Jones interactions at every timestep and updating PME forces every second timestep.

Antibody equilibration followed a 3-steps protocol: (1) 2000 steps of conjugate-gradient energy minimization, (2) a 200 ps NPT run with a temperature ramp from 0 K to 300 K, and (3) an additional 200 ps NPT simulation at 300 K. The final snapshot from stage (iii) provided starting coordinates and velocities for production trajectories of 30 ns per antibody. Trajectory analysis was carried out in VMD, extracting 50 frames from the last 24 ns of each simulation after recentering and aligning the protein within the simulation box.

### In silico physicochemical profiling of antibodies

Physicochemical descriptors were calculated from full-length antibody sequences and homology models. Expasy ProtParam (https://web.expasy.org/protparam/) was used to calculate the number of residues, molecular weight, isoelectric point, aliphatic index, and GRAVY (grand average of hydropathy) scores of the antibodies. Solvent-accessible surface areas (SASAs) were computed for categorized residues: hydrophobic (ALA, VAL, LEU, ILE, PRO, PHE, MET, TYR, TRP), aliphatic (ALA, VAL, LEU, ILE), aromatic (PHE, TYR, TRP), polar (SER, THR, ASN, GLN, TYR), positively charged (ARG, HIS, LYS), and negatively charged (ASP, GLU). Histidine protonation at pH 6.0 was modeled with 50% occupancy. Surface patches were identified using DBSCAN clustering of exposed residues (SASA > 0.1 nm^2^), with mean patch areas calculated for each residue category. Additional parameters were directly calculated from the MOE, including net charge, radius of gyration, sedimentation coefficient, dipole moment, affinity between light and heavy chains (affinity HC/LC), and buried surface area between light and heavy chains (BSA HC/LC). All structure-based properties were averaged across 50 frames to account for conformational dynamics of proteins in solution.

### Feature selection and clustering

Five feature clusters were considered including size, charge, polarity, hydrophobicity, and interfacial properties. Properties within the size and hydrodynamic cluster includes number of amino acid residues, molecular weight, and protein volume. Charge concerns isoelectric point (pI), net charge at pH 6, variable-domain Fv net charge, fraction of positively/negatively charged residues, and positively/negatively charged SASAs. Polarity concerns dipole moment and polar SASA. Hydrophobicity concerns aliphatic index, GRAVY score, and hydrophobic/aromatic/aliphatic SASAs. Interfacial properties concern affinity between HC/LC and buried surface area between HC/LC.

Features within each cluster were z-score normalized and physically aligned such that higher values uniformly represent increased property magnitude (e.g., higher Charge composite score indicates more positive character). Specifically, composite scores were calculated as equal-weighted averages of aligned z-scores, i.e., 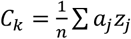, where *z*_*j*_ are z-scored feature *j* and a_*j*_ = ±1 align features to uniform directions. This yielded five interpretable metrics for each antibody.

## Author Contributions

Z.H., N.A.E., P.S., A.E. and T.P.J.K conceived the study. P.S. and T.P.J.K. supervised the project. Z.H. designed and performed the experimental work. Z.H., G.L., and P.S. performed the in silico modeling and analysis. A.E. provided the antibodies. O.P. assisted with protein purification. R.S. assisted with droplet image analysis. Z.H. wrote the manuscript with inputs and feedback from A.E., G.L., P.S., and T.P.J.K.

## Acknowledgements

We would like to acknowledge funding from the European Research Council under the European Union’s Seventh Horizon 2020 research and innovation program through the ERC grant DiProPhys (agreement ID 101001615, T.P.J.K.). P.S. is a Royal Society University Research Fellow (grant no. URF\R1\201461) and acknowledges funding from UK Research and Innovation (UKRI) Engineering and Physical Sciences Research Council (EPSRC grant no. EP/X024733/1). N.A.E. acknowledges the National Growth Fund Big Chemistry from the Ministry of Education, Culture and Science (1420578).

