## Supplementary Information for "Molecularly specific solubilization of therapeutic antibodies"

Zexiang Han *et al.*

**a**

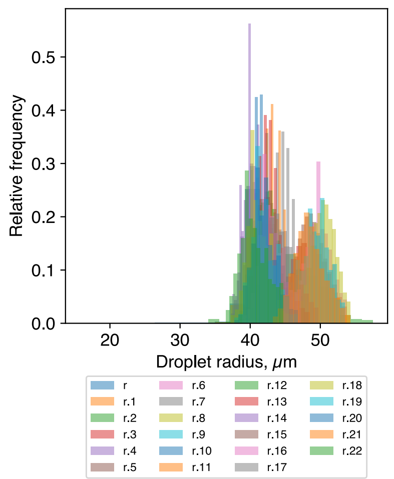


**b**

| Exp | # of droplets | Mean Radius | Std Dev | CV | PDI |
|---|---|---|---|---|---|
| r | 4962 | 40.89168 | 2.155305 | 0.052708 | 0.002778 |
| r.1 | 23199 | 41.88654 | 2.10961 | 0.050365 | 0.002537 |
| r.2 | 16491 | 46.95904 | 3.662045 | 0.077984 | 0.006081 |
| r.3 | 18607 | 48.3993 | 2.460855 | 0.050845 | 0.002585 |
| r.4 | 30752 | 40.03294 | 1.024978 | 0.025603 | 0.000656 |
| r.5 | 28731 | 48.61637 | 1.741607 | 0.035823 | 0.001283 |
| r.6 | 18067 | 49.73892 | 1.760711 | 0.035399 | 0.001253 |
| r.7 | 25257 | 44.60259 | 1.294132 | 0.029015 | 0.000842 |
| r.8 | 27320 | 50.50096 | 1.599977 | 0.031682 | 0.001004 |
| r.9 | 6731 | 49.23789 | 1.941091 | 0.039423 | 0.001554 |
| r.10 | 35647 | 41.40694 | 1.092558 | 0.026386 | 0.000696 |
| r.11 | 41549 | 43.15175 | 1.216182 | 0.028184 | 0.000794 |
| r.12 | 37863 | 42.2436 | 1.413235 | 0.033454 | 0.001119 |
| r.13 | 33567 | 42.51635 | 1.213694 | 0.028547 | 0.000815 |
| r.14 | 30028 | 40.63679 | 1.391851 | 0.034251 | 0.001173 |
| r.15 | 33459 | 42.90003 | 1.958699 | 0.045657 | 0.002085 |
| r.16 | 19353 | 43.08518 | 1.72495 | 0.040036 | 0.001603 |
| r.17 | 33948 | 44.26532 | 2.179097 | 0.049228 | 0.002423 |
| r.18 | 44776 | 41.10646 | 1.535368 | 0.037351 | 0.001395 |
| r.19 | 36606 | 41.92981 | 1.663297 | 0.039669 | 0.001574 |
| r.20 | 29039 | 42.02264 | 1.673515 | 0.039824 | 0.001586 |
| r.21 | 18757 | 48.4897 | 2.184624 | 0.045053 | 0.00203 |
| r.22 | 45418 | 40.51622 | 1.891151 | 0.046676 | 0.002179 |

**Supplementary Figure S1. Size distribution of water-in-oil microdroplets. a**, Apparent microdroplet size distribution histograms sampled from 23 representative microfluidic experiments. Note that while the mean droplet size can vary slightly between experiments, the measured phase boundary has been shown to be droplet volume-insensitive.[22] **b**, Table summarizing the number of droplets analyzed, apparent mean microdroplet radius in μm, standard deviation, coefficient of variation (CV) and polydispersity index (PDI).

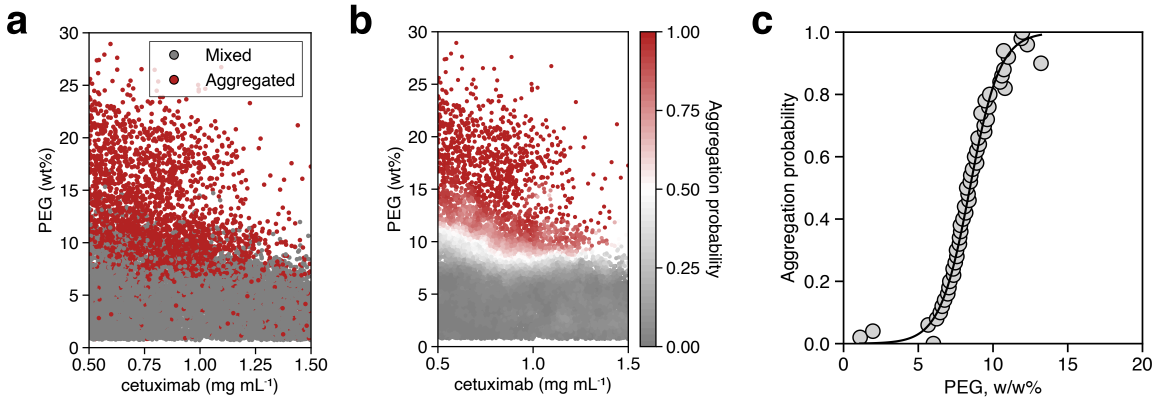


**Supplementary Figure S2. Determination of the solubility profile of an antibody. a**, Relative solubility phase diagram of cetuximab with binary classification of data points, as shown in Figure 1b. **b**, Averaged solubility phase diagram, where local aggregation probability was computed by spatial averaging of neighboring measurements within the composition space. The averaged representation highlights the underlying phase boundary while preserving the experimentally sampled composition range. **c**, Aggregation probabilities at 1 mg $mL^{-1}$ for various PEG concentrations were extrapolated from the phase diagram, and a sigmoidal fit was used to calculate antibody relative solubility as the PEG concentration for which the aggregation probability is 50%. Here, the relative solubility of cetuximab was determined to be 8.5±1.6 wt%.

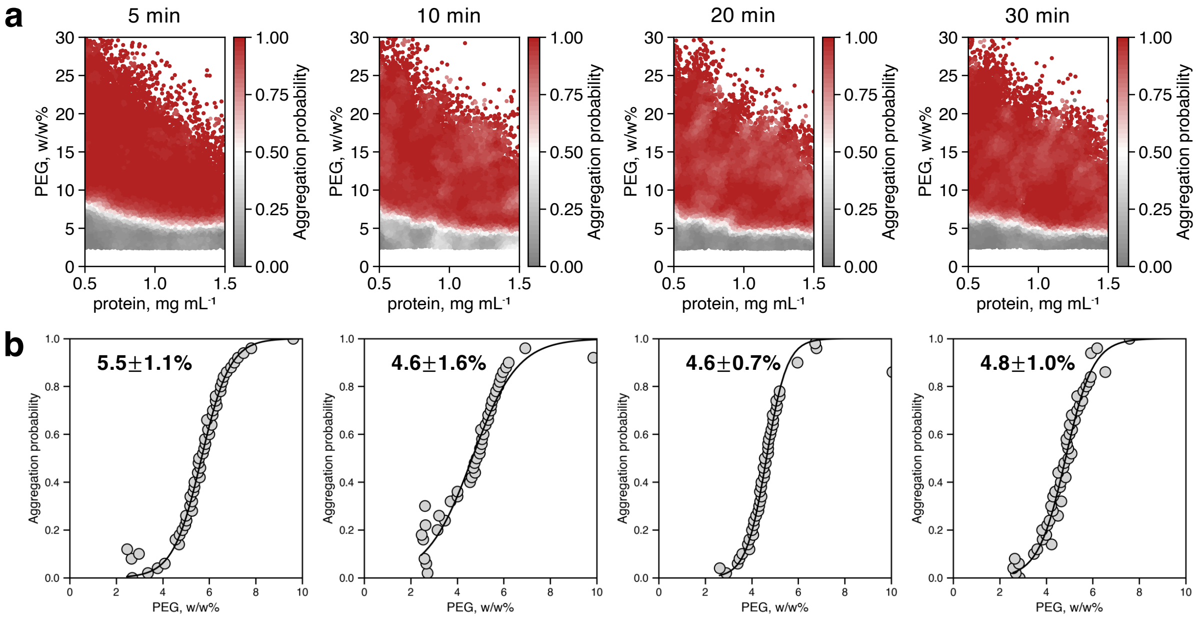


**Supplementary Figure S3. Effect of microdroplet incubation time on the measured antibody solubility. a,** Measured solubility phase diagrams of a model mouse IgG1 antibody following increasing durations of on-chip or on-line incubation. **b**, Corresponding sigmoidal fits and the determined relative solubilities at 1 mg mL$^{-1}$. The phase boundaries show insignificant variations with time. These results illustrate the robustness of the screening method, where changes in incubation time beyond 5 min minimally affect the solubility profile. This is likely generalizable to the IgG antibodies studied in this work.

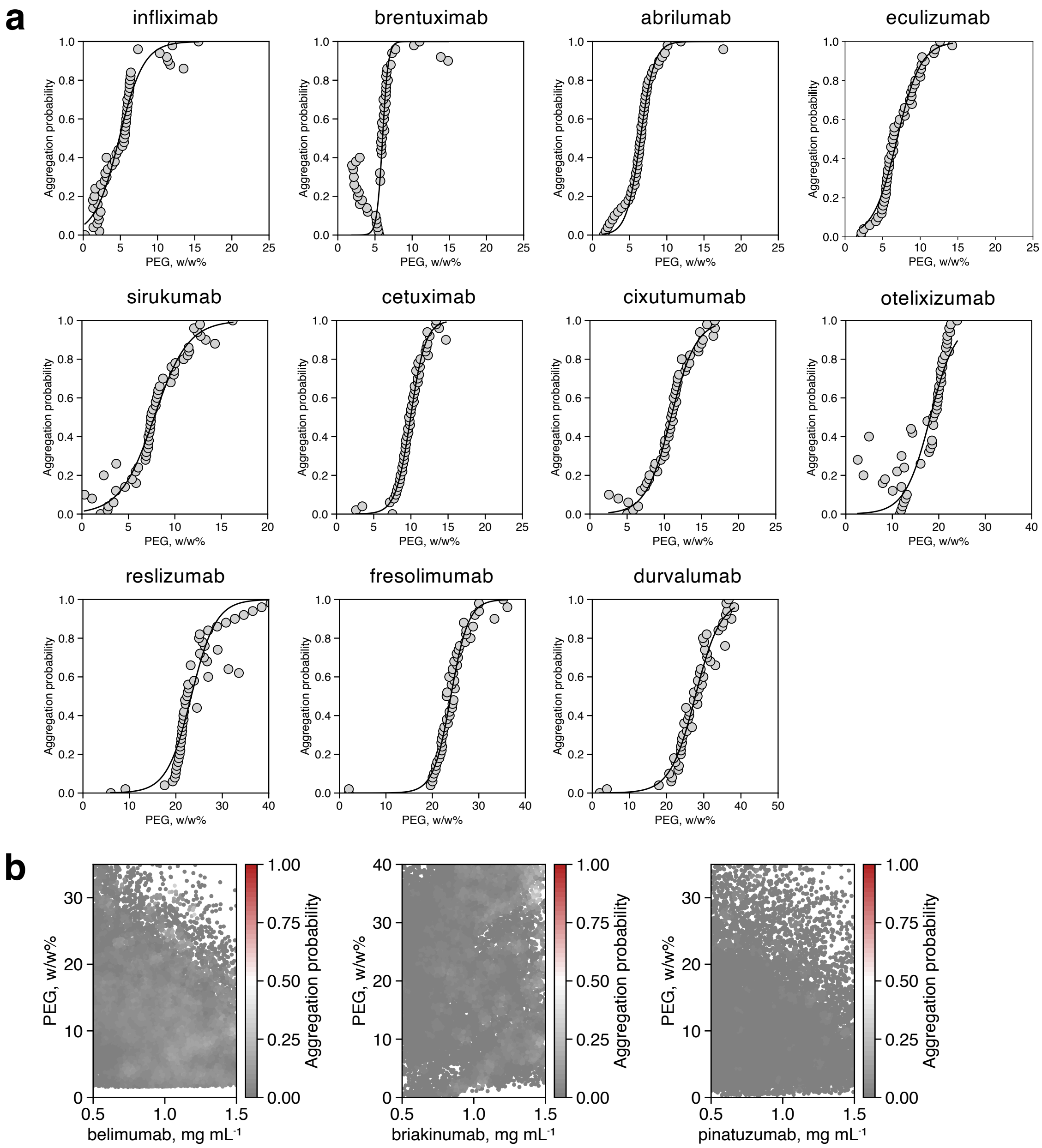


**Supplementary Figure S4. Measured solubility profiles of a panel of therapeutic antibodies.** **a**, Solubility profiles of 11 antibodies with sigmoidal fits. **b**, Phase diagrams of three mAbs (belimumab, briakinumab, pinatuzumab) that did not precipitate out under the assay conditions, showing only mixed phase.

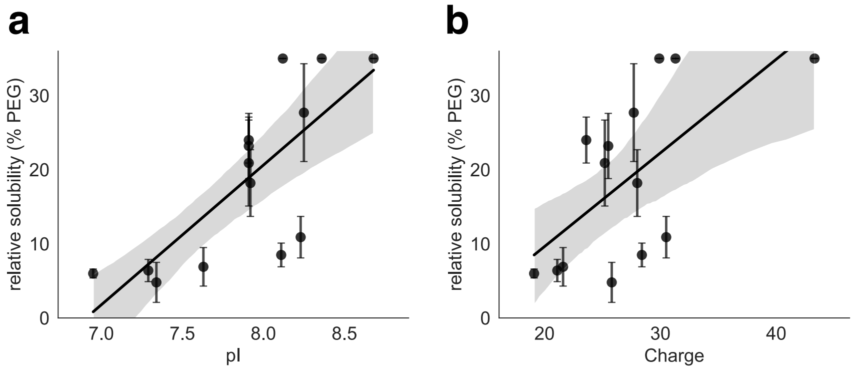


**Supplementary Figure S5. Correlation between mAb relative solubility and its charge. a,** Calculated full-length isoelectric point. **b,** Calculated charge at pH 6.0.

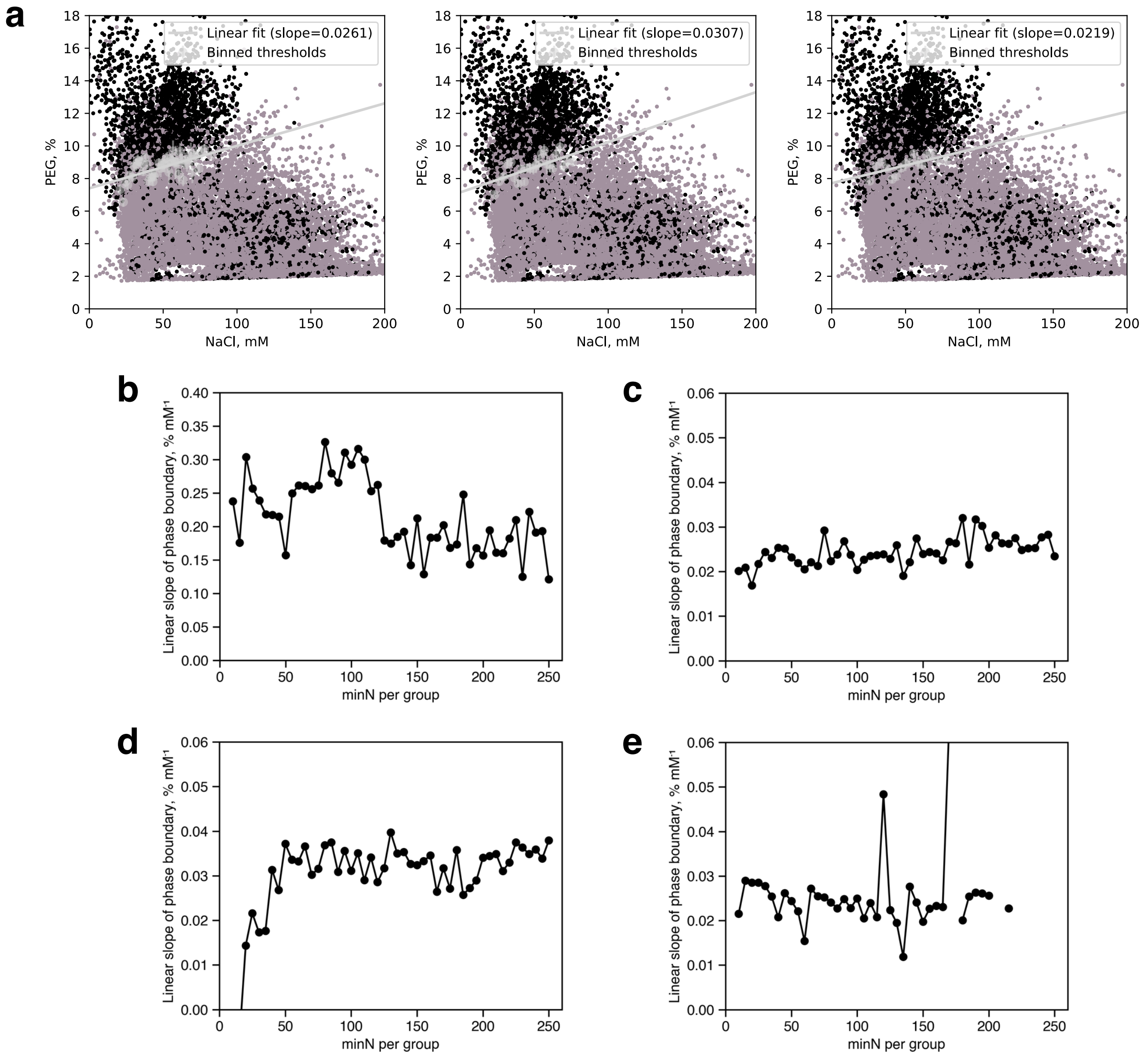


**Supplementary Figure S6. Calculation of solubilization scores. a,** Cetuximab-Arg phase diagrams showing the extrapolation of phase boundary slope using different binning size minN = 50, 100, and 150. **b-e,** Fitted slope as a function of the minimum cluster size applied to (b) His, (c) NaCl, (d) Arg, and (e) sucrose.

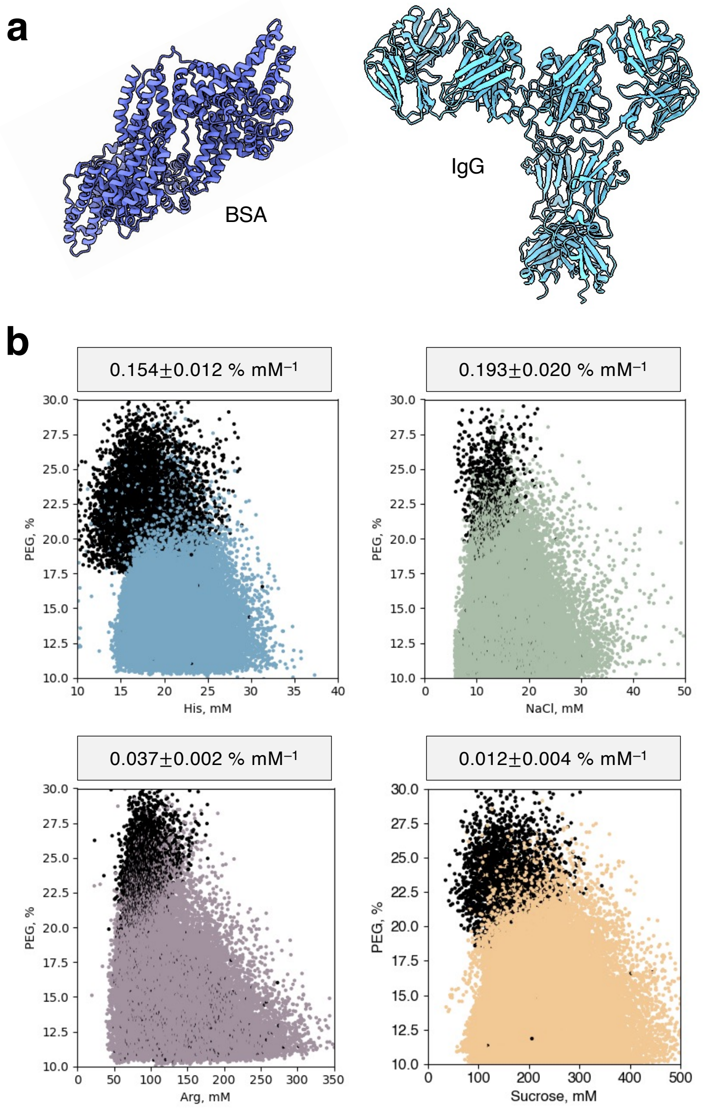


**Supplementary Figure S7. Excipient effects on bovine serum albumin (BSA) in standard formulation buffer. a,** Protein structure of BSA (PDB 3V03) compared to an IgG protein cetuximab. **b,** Solubility phase diagrams of BSA in the presence of histidine ($n$ = 36,163), sodium chloride ($n$ = 27,892), arginine ($n$ = 30,495), and sucrose ($n$ = 41,080) and determined solubilization scores.

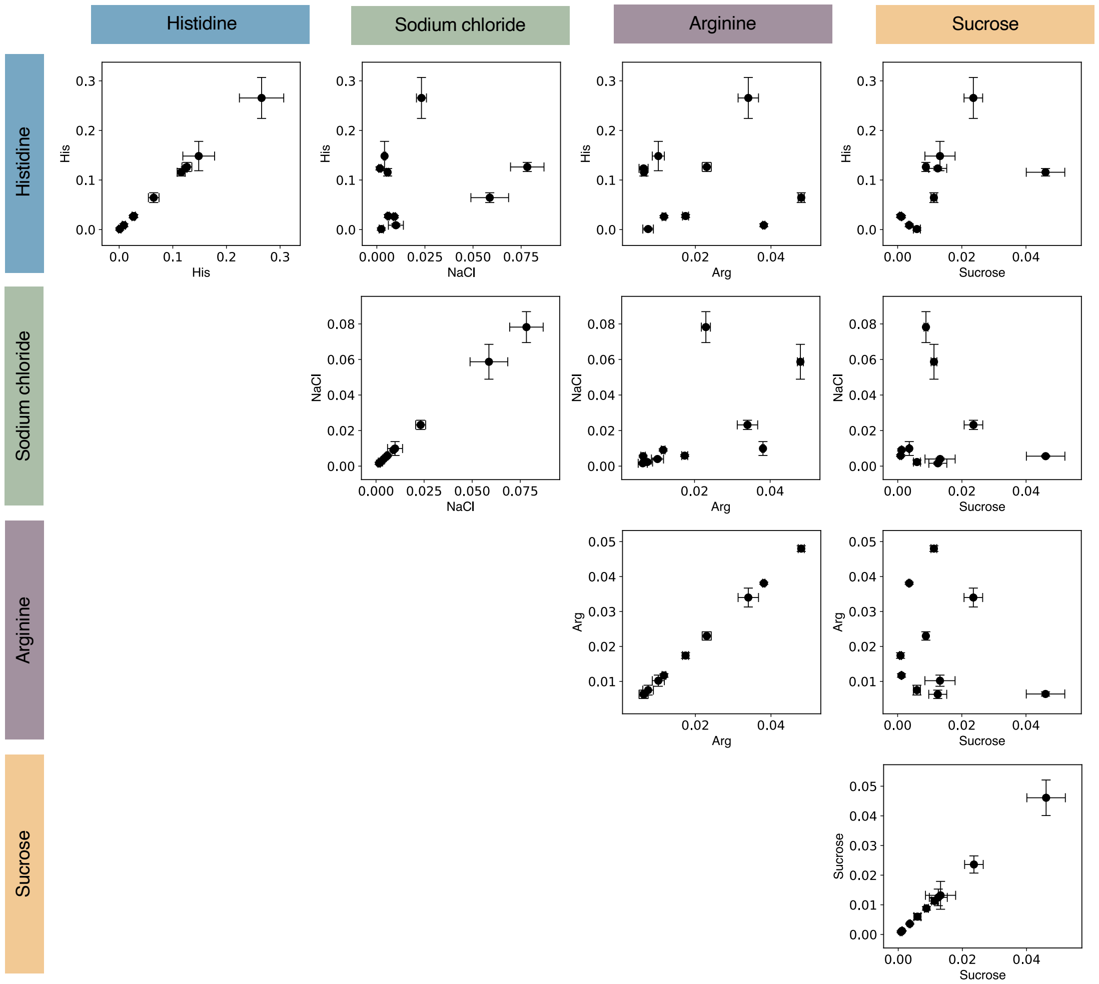


**Supplementary Figure S8. Scatterplots showing the cross-correlations of solubilization scores.** Units in % $mM^{-1}$.

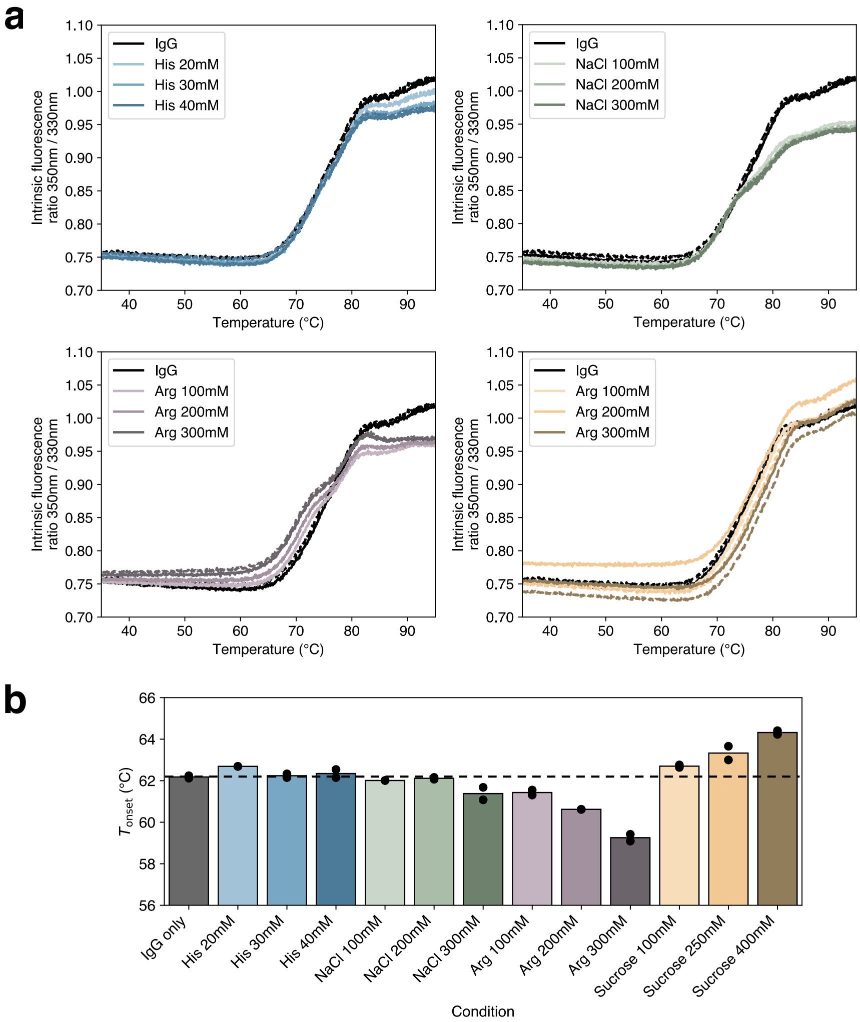


**Supplementary Figure S9. Thermostability analysis. a,** Thermal unfolding curves of cetuximab in the absence vs. presence of formulation excipients at different concentrations. Measurements were taken in duplicates for each condition, shown in solid and dashed lines, respectively. **b,** Effect of excipient addition on the onset temperature $T_{onset}$ of mAb.

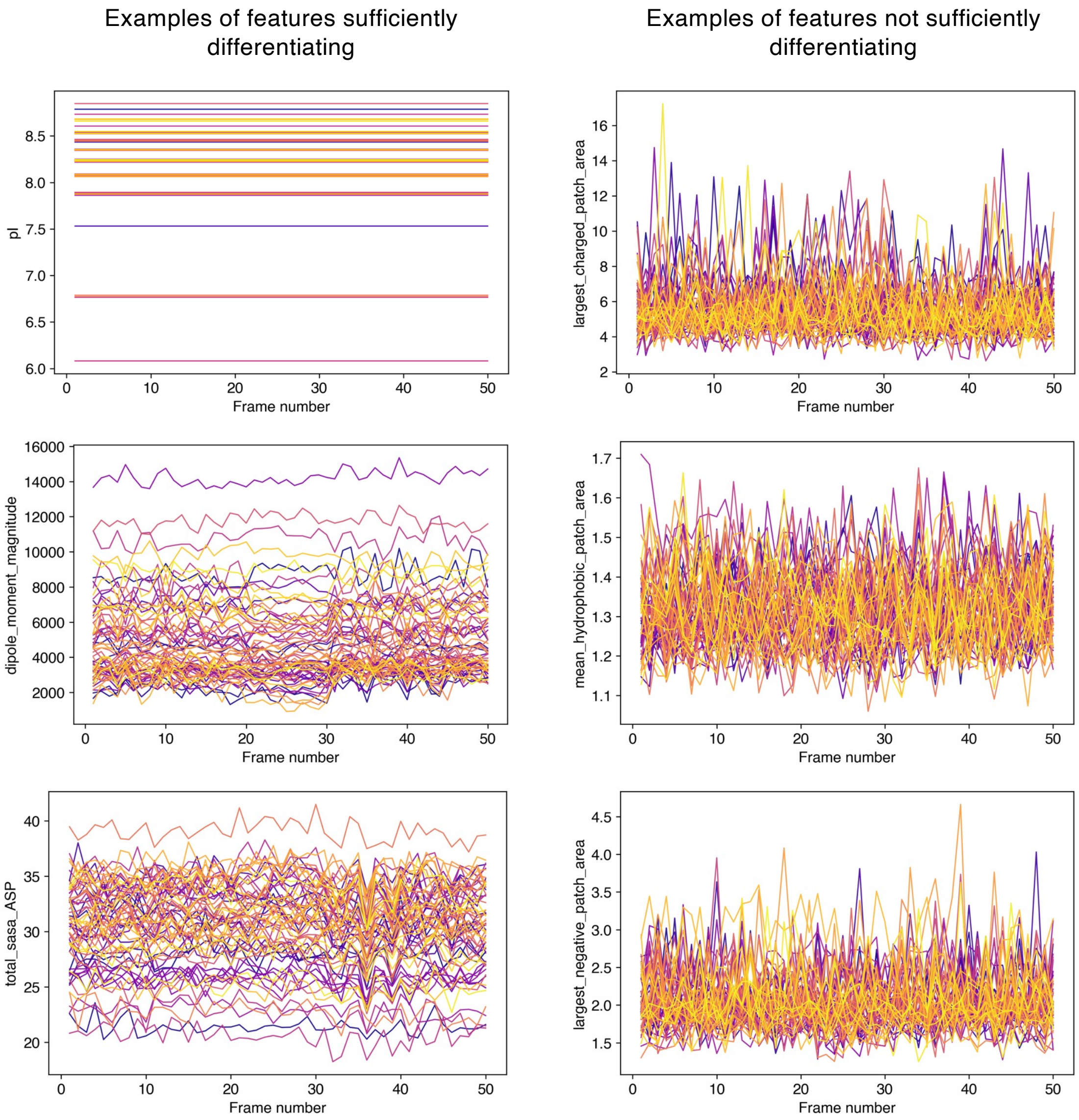


**Supplementary Figure S10**. **Identification of differentiating physicochemical features under conformational dynamics.** Features that are sufficiently differentiating across the antibody panel were retained for subsequent analysis. The remaining ones, notably patchiness-related metrics, were excluded.

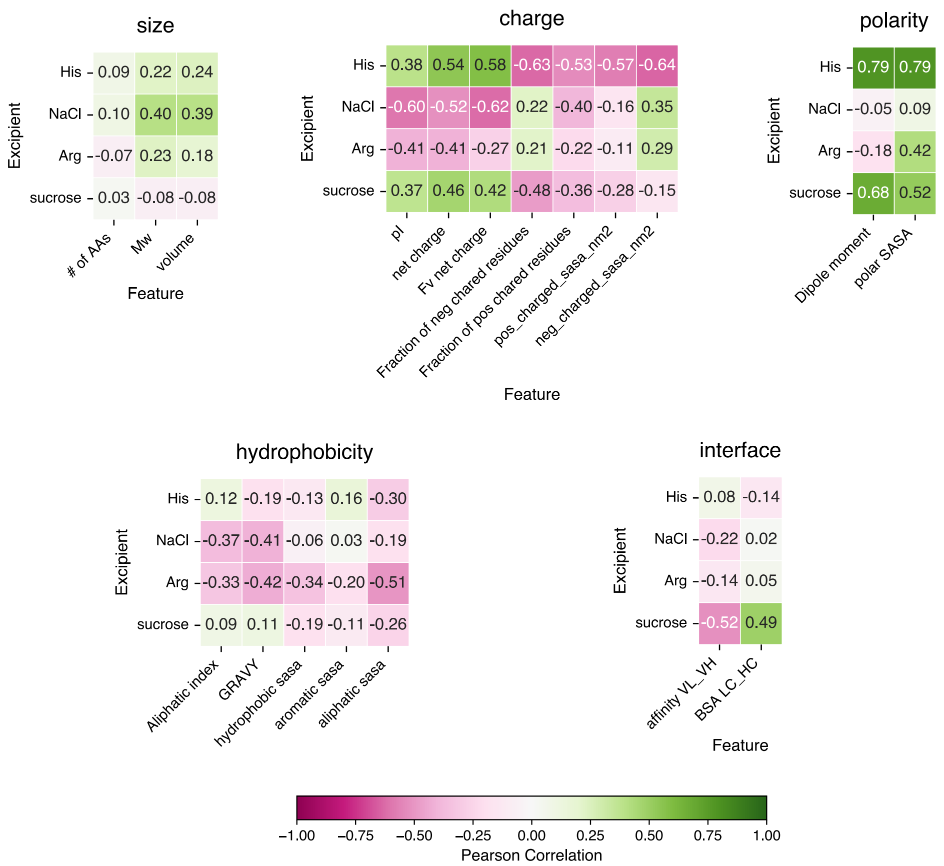


**Supplementary Figure S11. Correlation analysis against specific mAb molecular features.**
